# *linc2function*: A deep learning model to identify and assign function to long noncoding RNA (lncRNA)

**DOI:** 10.1101/2021.01.29.428785

**Authors:** Yashpal Ramakrishnaiah, Levin Kuhlmann, Sonika Tyagi

## Abstract

**Motivation:** LncRNAs are much more versatile and are involved in many regulatory roles inside the cell than previously believed. Existing databases lack consistencies in lncRNA annotations, and the functionality of over 95% of the known lncRNAs are yet to be established. LncRNA transcript identification involves discriminating them from their coding counterparts, which can be done with traditional experimental approaches, or via *in silico* methods. The later approach employs various computational algorithms, including machine learning classifiers to predict the lncRNA forming potential of a given transcript. Such approaches provide an economical and faster alternative to the experimental methods. Current *in silico* methods mainly use primary-sequence based features to build predictive models limiting their accuracy and robustness. Moreover, many of these tools make use of reference genome based features, in consequence making them unsuitable for non-model species. Hence, there is a need to comprehensively evaluate the efficacy of different predictive features to build computational models. Additionally, effective models will have to provide maximum prediction performance using the least number of features in a species-agnostic manner.

It is popularly known in the protein world that “structure is function”. This also applies to lncRNAs as their functional mechanisms are similar to those of proteins. Generally, lncRNA function by structurally binding to its target proteins or nucleic acid forming complexes. The secondary structures of the lncRNAs are modular providing interaction sites for their interactome made of DNA, RNA, and proteins. Through these interactions, they epigenetically regulate cellular biology, thereby forming a layer of genomic programming on top of the coding genes. We demonstrate that in addition to using transcript sequence, we can provide comprehensive functional annotation by collating their interactome and secondary structure information.

**Results:** Here, we evaluated an exhaustive list of sequence-based, secondary-structure, interactome, and physicochemical features for their ability to predict the lncRNA potential of a transcript. Based on our analysis, we built different machine learning models using optimum feature-set. We found our model to be on par or exceeding the execution of the state-of-the-art methods with AUC values of over 0.9 for a diverse collection of species tested. Finally, we built a pipeline called *linc2function* that provides the information necessary to functionally annotate a lncRNA conveniently in a single window.

**Availability:** The source code is accessible use under MIT license in standalone mode, and as a webserver (https://bioinformaticslab.erc.monash.edu/linc2function).

## 1 Introduction

LncRNAs account for 80% of RNA transcribed in the cell [1, 2]. These are emerging as master regulators of gene expression via various interaction mechanisms with other biomolecules at the transcription, translation, or epigenetic level [3]. These RNAs are involved in various cellular biological processes, and they do that in conjunction with other biomolecules through complex pathways [4]. A large number of lncRNAs are being discovered but about 95% of lncRNAs still lack functional annotation [5]. We reviewed existing lncRNA annotation methods and databases, some are sourced reliably while others are crowdsourced, and discovered that there is very little consensus between them [6]. Inconsistencies between databases is a major concern, and require newer methods to annotate these RNAs reliably. For instance, we found in our analysis that 230 genes were omitted from GENCODE human release v34, however, they are present in the previous version v24. Additionally, 41 of them are verified to play a role in various diseases as per the LncRNADisease 2.0 database [7, 8]. Furthermore, other than a few species with well-studied reference genomes, there is very little to no lncRNA annotations available for the remaining genomes. This demands reference-free or /textitab initio methods to study these RNAs.

Dysfunction of lncRNA can lead to cellular disequilibria such as in a disease state [9, 10]. Recent studies have revealed that many diseases are caused by misregulation or mutation of lncRNAs [10]. As an example, approximately 80% of experimentally validated lncRNAs reported in the LncBook database [11] are associated with more than 400 diseases; and more than 200,000 predicted disease associations are present in LncRNADisease 2.0 database [7, 8]. Therefore, understanding of functional mechanisms of lncRNA would help in the diagnosis, prognosis, prevention, and treatment of several disorders.

LncRNA functions by folding into a secondary (2D) and tertiary (3D) structure containing multiple structural domains [12, 13, 14]. Moreover, the binding target which are the biomolecules such as DNA, RNA, and proteins, is mainly determined by its 2D structure [15]. Consequently, in order to retain its function, the secondary structure of the lncRNAs is under a higher evolutionary pressure resulting in higher conservation than its sequence [16]. To confirm this hypothesis, we looked for ultra conserved regions (UCRs) around 65 lncRNA genes implicated in endometriosis, as sourced from the FANTOM-CAT dataset [17]. We did not find any overlap of UCR regions with the exonic loci of these genes indicating a poor sequence conservation. On the other hand, we observed 7 out of the 65 genes had UCR regions in non-exonic regions in this study. More details are available here: https://gitlab.com/tyagilab/linc2function/-/blob/master/HumanDiseaseUcr/human_disease_ucr.md.

Thus, determining lncRNA folding and its structural domains are crucial in unlocking its function in disease aetiology, and determining its interactome [18, 19]. Any changes in the structure of lncRNA will result in a partial or complete breakdown of its functional pathways and result in cellular malfunction.

Some of the known lncRNA-DNA interactions involve forming triplex structures such as R-loops [20], and triple helices [21] that in turn are involved in transcriptional and post-transcriptional regulation, chromatin remodeling, and DNA repair [22]. Thus, a transcript with high Triplex Forming Potential (TFP) is more likely to be involved in these mechanisms either in *cis* or in *trans*.

RNA Binding Proteins (RBPs) form RNA-protein complexes by binding to lncRNAs and these complexes are associated with regulating various cellular pathways [23]. The functions of RBPs include transcription and translation regulation, DNA repair, splicing, apoptosis, and mediating stress responses [24]. By knowing the potential RBPs interactions with a lncRNA, it should be possible to estimate the functional pathways in which the lncRNA is involved, and these RBP interactions can be estimated by the presence of sequence or structural motifs on it.

LncRNA also interacts with other ncRNA and mRNA to perform their function [25]. LncRNA-miRNA interaction is a well studied phenomenon resulting in lncRNAs directly regulating the expression levels of a particular gene. MiRNAs can down-regulate the expression of their target mRNAs by binding in their UTR region [26]. LncRNAs can act as competing endogenous RNA (ceRNA) where they compete for miRNA binding along with mRNAs, as a result acting as regulators of corresponding mRNA targets [27, 28]. Thus, knowing the possible miRNA interactions of a lncRNA will enable us to know the genes that the lncRNAs can possibly modulate.

Accordingly, functional annotation of a lncRNA would consist of two parts: 1) distinguishing a lncRNA transcript from other non-coding and coding transcripts; and 2) identifying its functional structure along with its interacting biomolecules and their contact sites. We will refer to the set of interacting biomolecules as the lncRNA interactome.

Primary data to identify lncRNA comes from High throughput sequencing (HTS) experiments [29]. Short sequencing reads obtained from a HTS run are assembled to build RNA sequences. The assembled sequences are further analyzed to annotate them as various types of RNA transcripts. It is challenging to obtain features of a transcript that can be used to distinguish a lncRNA from other RNA transcripts due to their high diversity, and lack of comprehensive high-confidence reference data. However, a deep learning model can help identify such generic features in an *ab-initio* manner. When trained on existing known lncRNAs examples it can pick up similar patterns on screening the transcripts obtained in RNA-seq experiments *en masse*[30, 31].

Experimental methods to identify RNA interactome are expensive, time-consuming, and cannot be performed for each cell type or state. Moreover, the expression level of these lncRNAs is very low compared to mRNAs and also tissue-specific in nature, which pose further challenges for *in vitro* methods. Hence, *in silico* approaches are a favourable choice to predict the lncRNA interactome.

Many *in silico* tools are developed to identify if a given sequence is lncRNA using different machine learning algorithms [32, 33, 34]. We recently reviewed existing methods and current challenges in the process [6]. Some of these methods are reference-based in which case a given transcript sequence is compared against a pre-annotated reference for annotation purpose. Other methods are *ab initio* that do not require a reference, and predictions are made based on primary and derived properties of a transcript sequence. Reference-based methods are limited in their ability to perform cross-species predictions, and therefore, can not be applied for non-model organisms lacking a pre-annotated reference. Structural attributes are shown to be more important than its sequence, and show higher evolutionary conservation [15]. Further, identification of structural domains and interactome are required for full functional annotation of lncRNA, but existing tools rarely make use of these characteristics (refer Table 1).

**Table 1.**
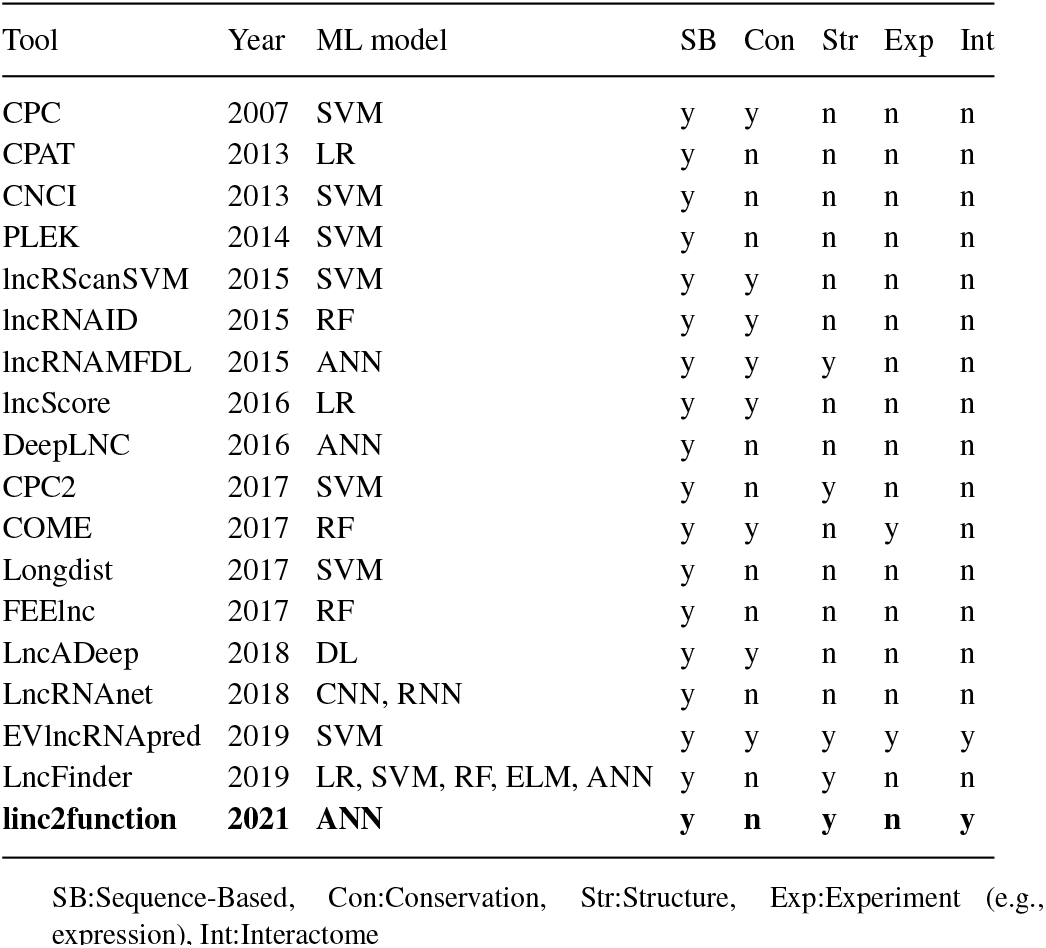
Table listing different tools, year of release, machine learning model used, and the type of features used (Sequence-Based, Conservation, Structure, Experiment (e.g., expression), and Interactome. Row containing linc2function is highlighed in bold text)

To summarise, in this study first, we aim to develop a machine learning model to identify lncRNA in a species-agnostic manner; and secondly, to build a comprehensive pipeline for annotating lncRNA.

We propose an artificial neural network (ANN) that can identify the lncRNA forming potential of a given transcript in a species-agnostic manner. Using non-reference features extracted and diligently selected from its sequence, structure, and interactome characteristics. We extend this model to build a pipeline called *linc2function*, containing an identification module, structural, and interactome modules by integrating it with existing open-source methods for structural and interactome predictions. It is deployed both as a web-server, and a standalone tool under the MIT license for wider accessibility. To the best of our knowledge, such a pipeline does not exist, and we believe it will be of great significance to advance lncRNA research for biomedical applications.

## 2 Methods

### 2.1 lncRNA identification

#### 2.1.1 Data Source

Various open-source repositories are available online that contain data related to lncRNAs. Some of them contain only the identified lncRNAs and their sequences, whereas others augment their structures, functions, and disease associations. The information contained in these repositories was obtained by manually curating the literature, low-throughput experiments, high-throughput experiments, *in silico* predictions, or a combination of any of these. Further, the information gathering might be handled by domain experts, crowdsourced, or generated via computer automation. Besides their varied origin, there are complications in comparing them directly as the identifiers used by each one of these repositories are different [6]. For the purpose of obtaining consensus data from all four repositories we opted to use the genomic coordinates matching with a tolerance of plus or minus five nucleotide positions. This consensus data as depicted in Fig 1 was used as positive training set for our ANN model. An equal number of coding sequences were selected randomly from the GENCODE database.

**Fig. 1.**
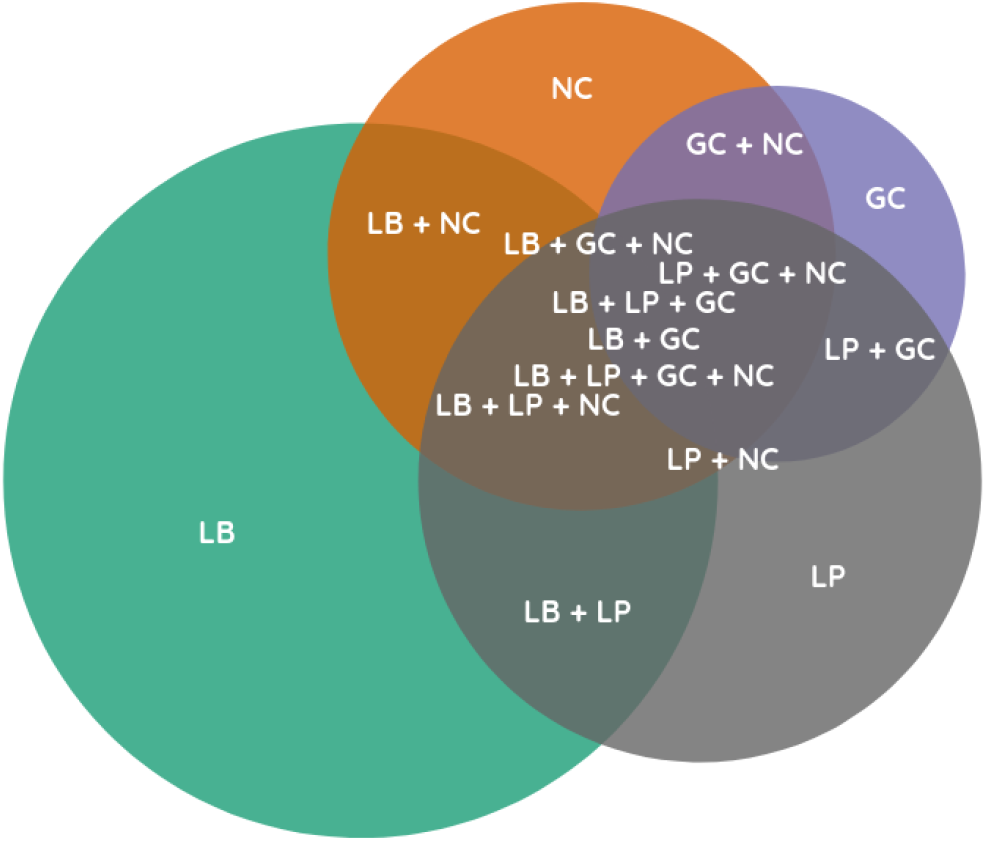
Venn diagram showing the overlap of the transcripts between LncBook (blue), LNCipedia (orange), GENCODE (green), and NONCODE (red). In the figure LB refers to the LncBook, LP refers to the LNCipedia, GC refers to the GENCODE and NC refers to the NONCODE

#### 2.1.2 Feature Extraction

Various sequence-based, structural, and interactome features are extracted from the curated nucleotide sequence data as described in section 2.1.1. We analyzed the effectiveness of each feature in predicting lncRNA forming potential in a standalone manner as well as in conjunction with other features. For this purpose, we collected as many features as practically possible. A hierarchical tree diagram of the collected features is shown in Fig 2.

**Fig. 2.**
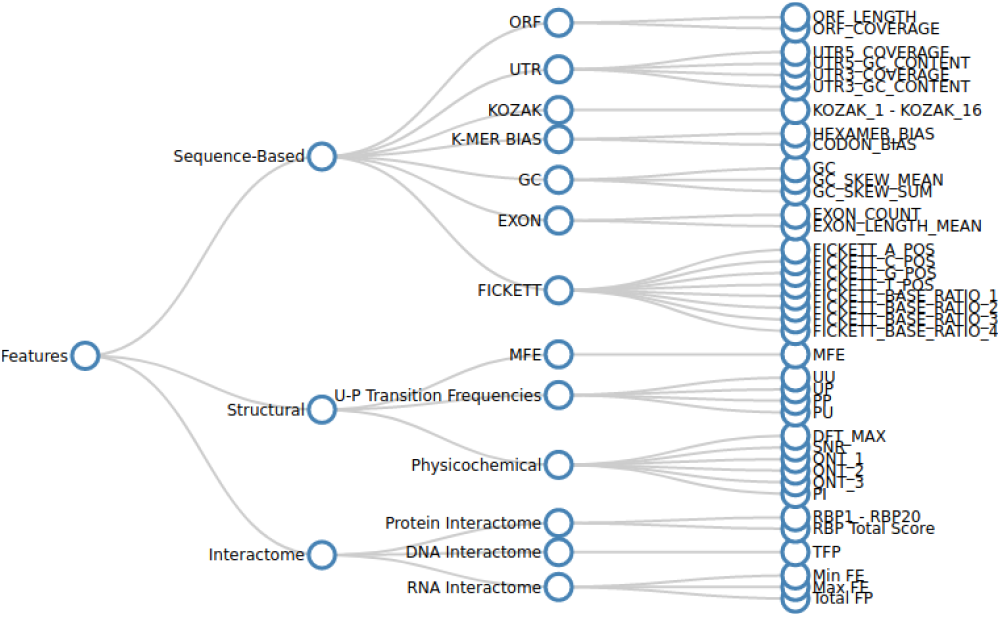
A hierarchical tree diagram illustrating different features and their categories considered in this study

Sequence-based features are obtained by the sequential characteristics of the transcript which quantifies its various nucleotide distribution patterns. Structural features are related to the properties of the secondary structure of a transcript, such as minimum free energy (MFE), paired-unpaired transition frequencies, and physicochemical features. Interactome features are the measure of nature and magnitude of the lncRNA interactions with their target RNA, DNA, or protein biomolecules.

#### 2.1.3 Feature Selection

In the first step, we eliminated the features using co-variance analysis, univariate feature selection techniques [35], and by feature importance measures using forests of trees. We looked at their co-variance, F-Value, Chi-Squared, Mutual Information, and feature importance measures from various tree-based classifiers. Primarily, this step is necessary to eliminate the features which contribute more noise quotient to the model than their contribution towards model prediction. In consequence, 35 features were retained after the exclusion.

Next, the recursive feature elimination (RFE) method is used to select the subset of features that contribute the most to the discriminative ability of the model. RFE computes feature importance by building a linear model and calculating the coefficients. Then the least important features are pruned recursively until a predetermined stopping condition is met as shown in Fig 3. Having fewer features also results in reduced dimensionality of the input feature set and therefore, lowering the training and prediction time. Obtaining some of the features can be a high compute intensive and time consuming proposition and rendering it impossible to screen large amounts of transcripts in real-time. Thus, we obtained the *feature_importances_attribute* value to be 10 to make a set of human-specific and species-agnostic features.

**Fig. 3.**
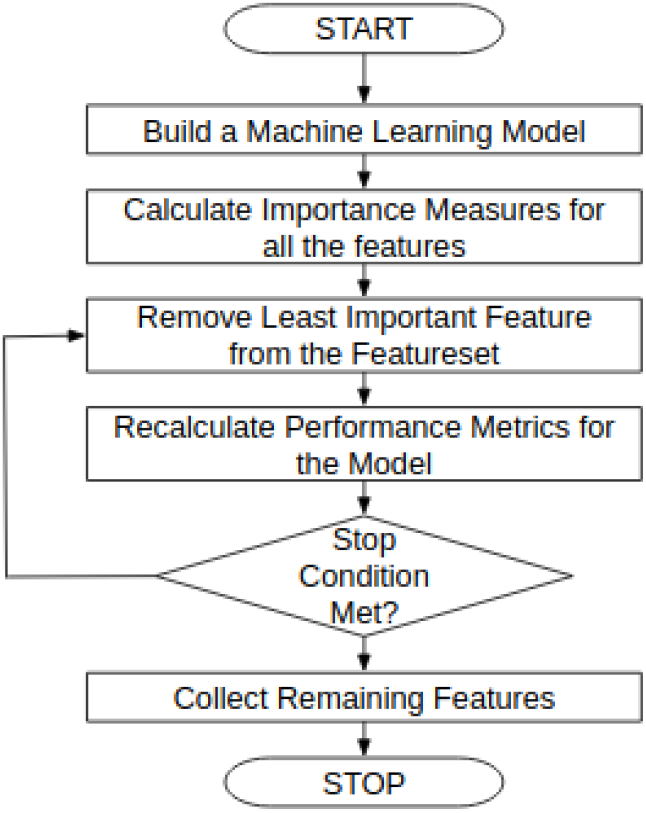
A flowchart explaining the steps involved in the RFE algorithm in detail

Thus,from the above steps, we create a “Full” feature set (n=35) and a “Light Weight” feature set (n=10). The model based on the former feature set would be computationally intensive as compared to the later. For constructing the models which are good at cross-species generalization, we shortlisted two types of features, one including the reference-based features, and the other one independent of the reference genome. By taking different combinations of feature sets and types, 4 models namely were build. As shown in Table 2, the models are called: HSF - Human Specific Full, HSLW - Human Specific Light Weight, SAF - Species Agnostic Full, and SALW - Species Agnostic Light Weight.

**Table 2.** A table containing different ANN models their acronym, number of features, and presence or absence of reference-based features)

|  | Acronym | Features | Reference Based Features |
| --- | --- | --- | --- |
| Species Agnostic Light Weight | SAL | 10 | Absent |
| Species Agnostic Full | SAF | 35 | Absent |
| Human Specific Light Weight | HSL | 10 | Present |
| Human Specific Full | HSF | 35 | Present |

SB:Sequence-Based, Con:Conservation, Str:Structure, Exp:Experiment (e.g., expression), Int:Interactome

#### 2.1.4 Model Architecture

We built four ANN models as listed in Table 2, and trained them with the shortlisted feature sets. The primary objective is to keep the complexity of the models as low as possible so that they are easily generalizable. The models consist of 1 input layer with the number of nodes equal to the number of input feature-length. The input layer is followed by a hidden layer with l1_l2 regularization to avoid overfitting and then a relu activation function (Fig 4). The l1_l2 regularization is applied to the kernel, bias, and activity values for this layer. There is a dropout layer of ratio 0.3 which randomly drops the nodes in the specified ratio during training to make the model more robust, and to improve model generalization. Finally, there is an output layer with one node and a sigmoid activation function that will provide the lncRNA forming potential for the given sequence. Setting an appropriate threshold for the lncRNA forming potential enables us to predict the two classes, positive class consisting of *lncRNA* and a negative class consisting of *not lncRNA*.

**Fig. 4.**
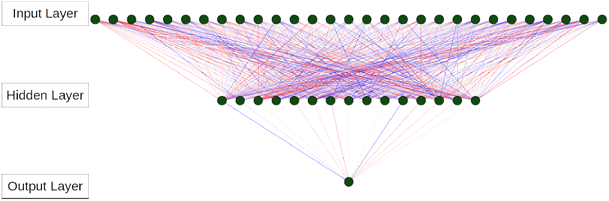
A representative neural network architecture for the ANN model showing input, hidden, and output layers. All the layers are densely connected with the color intensity of edges proportional to the connecting weights between two nodes. The negative edges are shown in red color and positive edges in blue color. The width of the edges is proportional to their weight.

#### 2.1.5 Model Training and Validation

Using our ANN architecture as explained in the previous section, we built the models using the python library Keras which is configured to use the TensorFlow library in the backend. We have trained the model using 80% of the observations present in data and the rest 20% was held out for validation.

Later, the models are used for prediction, where the classes (coding and noncoding) of the validation data set are predicted and compared against the true labels. Various classification metrics like Accuracy, Balanced Accuracy, Precision, Recall, F1-Score, and Area Under Curve (AUC) are calculated to establish the performance of the model. We also calculated the confusion matrix, which provides True Positives (TP), True Negatives (TN), False Positives (FP), and False Negatives (FN) using which we obtained Accuracy, Balanced Accuracy, Precision, Recall, and F1-Score.

#### 2.1.6 Cross-Species Comparison

One of the main objectives of this work is to build a model that can predict lncRNA given a transcript sequence regardless of the species. In this regard, we built two types of models, human-specific (HSLW and HSF) and species-agnostic (SALW and SAF). To test the cross-species predictions of our models, we shortlisted 8 species consisting of human, mouse, zebrafish, fruit fly, roundworm, yeast, wheat, and sea vase transcripts covering a wide evolutionary spectrum. The human and mouse data was obtained from the GENCODE [36] database and all the other species data was taken from the Ensembl [37] repository.

#### 2.1.7 Benchmarking

We benchmarked our methods against two other prominent tools currently available specifically, LncADeep [32] and LncRNAnet [34]. Both of these tools use neural networks in different configurations. LncAdeep uses an ANN built on a number of features covering sequence-based and conservation characteristics of a given sequence. LncRNAnet employs CNN and RNN which require only the sequence information which gets one-hot encoded of a fixed length. The benchmarking data consisted of randomly selected human transcripts from GENCODE, and randomly selected zebrafish and wheat transcripts from Ensembl databses.

### 2.2 lncRNA annotation

In this section we describe how we obtain interactome for the sequence provided.

LncRNA:DNA interactions were measured as the triplex forming potential (TFP). This was obtained by TriplexFPP [38], a deep learning-based utility. Similarly, for lncRNA:protein pairs we selected a list of RNA binding proteins (RBP) that are known to interact with lncRNAs, and used RBPDB [39] to scan sequences for RBP-binding sites. The RBPDB is a freely available database of RNA-binding specificities, and it is a collection of experimental observations of RNA-binding sites, both *in vitro* and *in vivo*, manually curated from the primary literature. Comprehensive lncRNA:RNA interactome analysis included interactions of lncRNA with mRNA, miRNA and other ncRNA. RIblast [40] was used to determine these RNA-RNA interactions, providing us an insight into the regulatory mechanisms in which the lncRNA might be involved. RIblast software is based on a seed-and-extension algorithm. We indexed ncRNA sequences from RNAcentral [41] and miRBase database [42] using RIblast db command to obtain RIblast database files. This indexed database is then used to scan for possible lncRNA-RNA interactions using RIblast search (ris) utility. The function of a lncRNA is determined by its structure, and its sequence plays very little role. We optimize an existing pipeline called SPOT-RNA [43] to predict an ensemble of two-dimensional structures in real-time by limiting the consensus models used within. We extended the length of sequences used in training and scanning.

### 2.3 The *linc2function* Pipeline

The above identification and annotation sub-modules were integrated to build the *linc2function* pipeline. The pipeline provides multiple characteristics of a lncRNA for the given sequence such as, the lncRNA forming potential of a given sequence followed by the prediction of its 2D structure, TFP, and its interactome. The pipeline is deployed in a web-server and as a standalone tool for wider accessibility.

## 3 Results

### 3.1 lncRNA identification

#### 3.1.1 Feature Engineering

An extensive set of features are extracted primarily to study the effect of an individual feature on predicting the lncRNA forming potential of a transcript. In order to build a machine learning model, there is a need to eliminate uninformative features. Although the machine learning algorithm which we choose (i.e. ANN) is inherently capable of performing feature selection by assigning less importance (weights) to the superfluous features, computing those avoidable features is a complex and time-consuming effort. To get an idea of which features are correlated with the target variable we plotted a heatmap as shown in the figure (Fig 5). It is observed from the heatmap that RBP and KOZAK motif related features have the least correlation, while all others have a noticeable relationship either positive or negative. The same observation was also confirmed by the feature selection process followed by us, where these features are usually ranked at the bottom. On the flip side, 5 out of the top 10 ranked features in our analysis are secondary structure based. Details of our feature importance analysis can be found in the Fig A1, Fig A2, Fig A3, Fig A4. This is in accordance with the studies showing that the secondary structures formed by the folding of the lncRNAs form the functional domains, through which they perform their biological functions. Additionally, the ORF features i.e ORF_LENGTH and ORF_COVERAGE are ranked highly along with EXON_COUNT reinforcing the importance of the presence or absence of ORF and exons in determining the lncRNA forming potential of a transcript. Hence, we decided to eliminate RBP and KOZAK motif related features from the initial cohort. Two full models HSF and SAF are built with human specific feature set containing all the remaining features, and species agnostic feature set containing non-reference features.

**Fig. 5.**
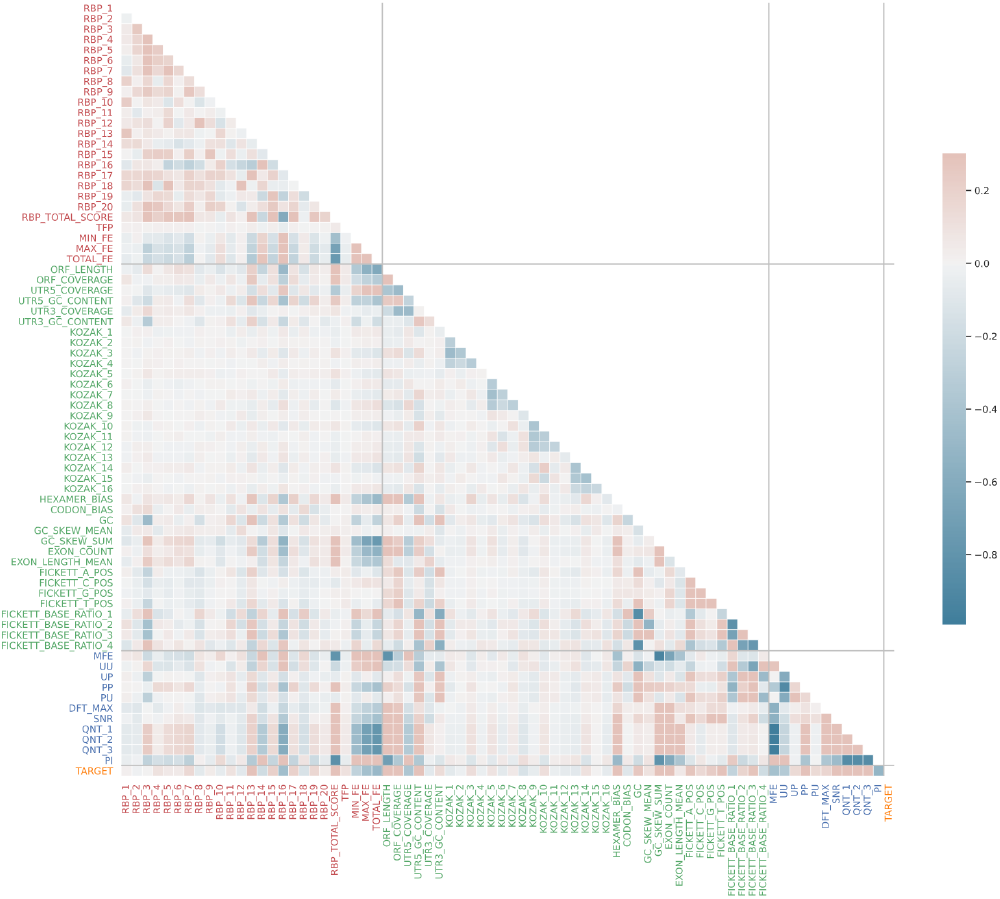
A heatmap containing all the feature and the target considered in this study. RBP-based features are labeled in red color, sequence-based features in green color, physicochemical features in blue, and the target in orange color. The color intensity of each cell corresponding to two features indicates the degree of correlation. Also, blue indicates a positive correlation and red indicates a negative correlation.

The HSF and SAF models contain certain features such as MFE, MIN_FE, MAX_FE, and TOTAL_FE that consume significantly more compute resources and time to calculate. Although, it will not be a problem in doing real-time analysis of individual transcripts, it is impractical to screen thousands of transcripts at once. Hence, we performed recursive feature elimination (RFE) where we found out only about 10 features are sufficient to get maximum prediction performance as shown in the figure 6. With this insight, we obtained the list of the top 10 human-specific and species agnostic features to build two light weight models i.e. HSLW and SALW.

**Fig. 6.**
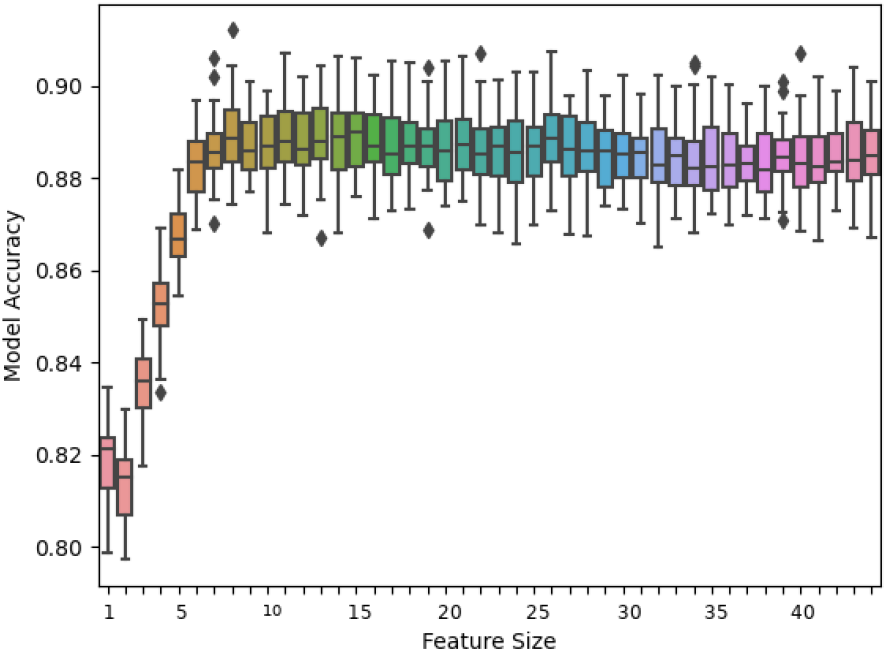
Cross-Validation Accuracies of the model on the training data for different values of n_features_to_select obtained by running RFE algorithm.

#### 3.1.2 Performance on human transcripts

As anticipated, when validated on unseen human transcripts, the human specific model performs marginally better than the corresponding species agnostic model. The decision to relinquish a small fraction of performance is taken with the aim of generalizing the model for cross-species predictions. Prediction probability distribution (Fig 7) shows a good separation between the classes. It can also be observed that models perform identically in predicting coding transcripts. However, the performance for noncoding transcripts varies slightly with HSF giving the best accuracy, followed by SAF, HSLW, and SALW in that order.

**Fig. 7.**
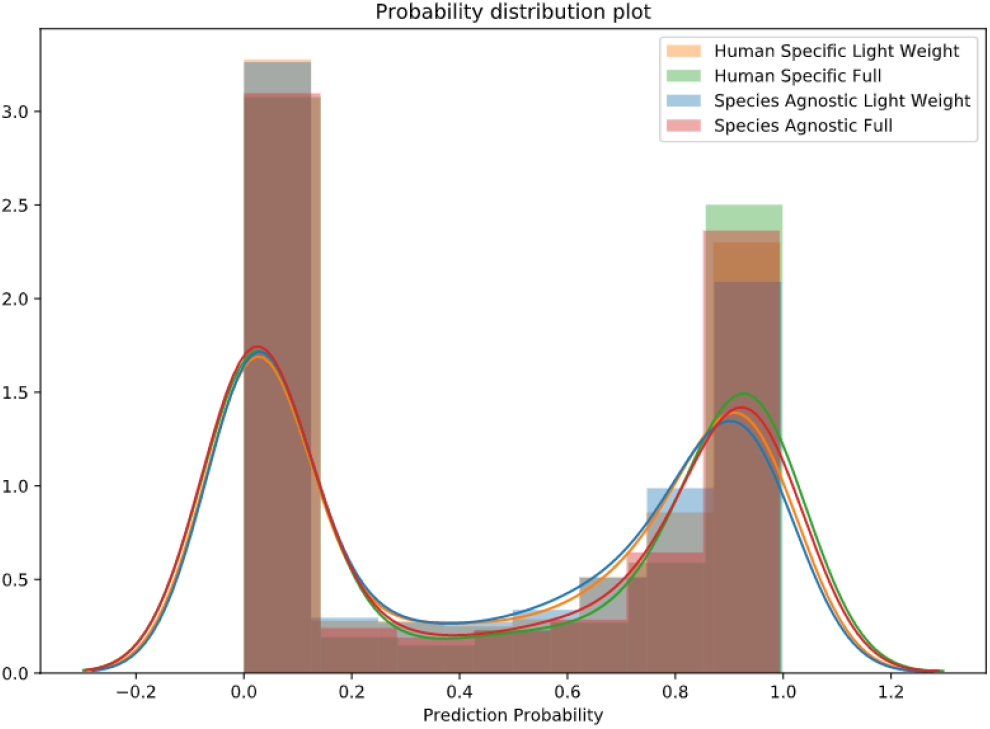
Histogram and distribution of predictions of both the models on randomly selected Human transcripts from Gencode. Blue - Human Specific. Orange - Species Agnostic. Predictions closer to 1.0 belong to noncoding will be labeled as noncoding and the ones closer to 0.0 as coding.

The testing ROC curve is drawn for both the models and obtained a graph as shown in Fig 8. From this figure, once more it can be observed that the human specific model is giving better predictions on the human dataset in comparison to the corresponding species agnostic model by a small margin. From the ROC curve, we calculated the AUC for the HSF model to be 0.9580, the HSLW model to be 0.9460, the SAF model to be 0.9528, and for the SALW model to be 0.9430. Overall, the models show AUC of above 0.94 on human transcripts with HSF performing marginally better than the others. Please refer appendix tables A1-A8 for more details on model performance on testing data.

**Fig. 8.**
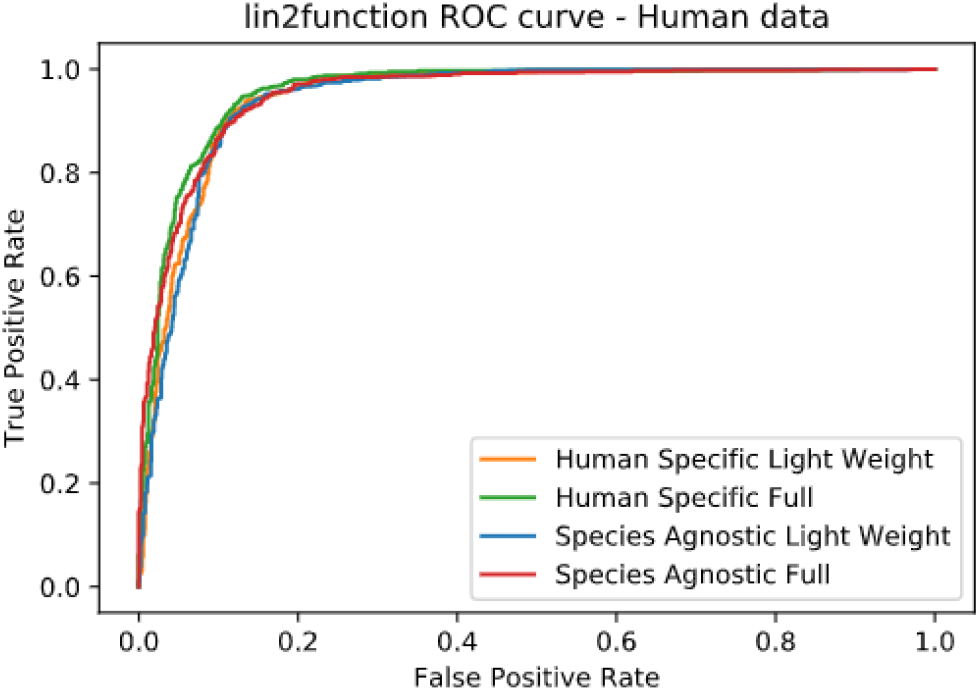
Testing ROC curve for linc2function Human Specific and Species Agnostic models evaluated on randomly selected Human transcripts from Gencode.

#### 3.1.3 Cross-species performance

It is observed that both the models are able to consistently achieve AUC of over 0.94 over different species as shown in Fig 9. As expected the HS model performed well on mammalian transcripts i.e. human and mouse, but on all the other species including vertebrates such as, zebrafish, the SA models prediction performance is superior. This demonstrates the SA model’s capability of performing well on other divergent species by capturing the generic characteristics of lncRNA transcript across species.

**Fig. 9.**
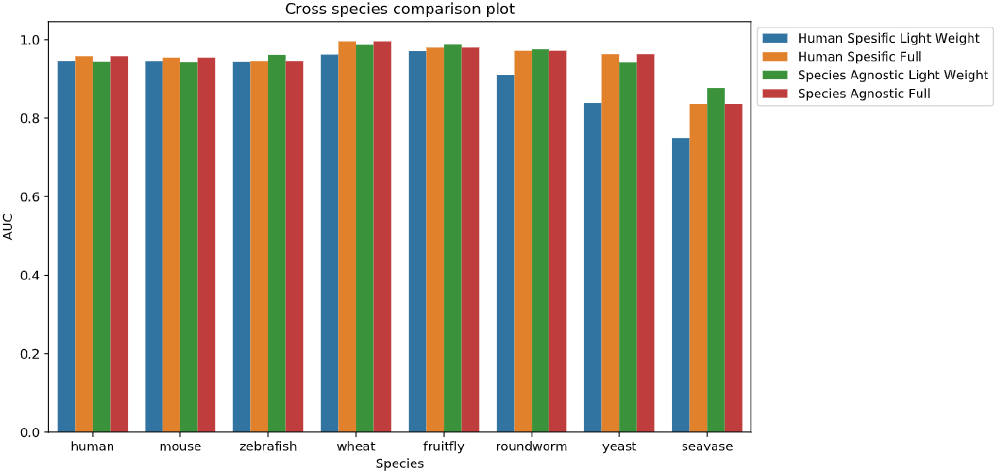
Cross-Species prediction performance comparing Balanced Accuracy Human Specific and Species Agnostic models evaluated on 8 different species (Fruit Fly, Human, Mouse, Roundworm, Sea Vase, Wheat, Yeast, Zebrafish in no particular order).

#### 3.1.4 Benchmarking

Benchmarking results (Fig 10, Fig 11, Fig 12) also show that the *linc2function* model is able to generalize well for species other than the one it was trained on. Its performance metrics are better for zebrafish dataset as against the human dataset when compared to the two state of the art methods namely, LncADeep [32] and LncRNAnet [34]. Moreover, its performance for the wheat dataset is as good as LncRNAnet and outperformed LncAdeep. Overall, all three models were able to consistently achieve an AUC value of around 0.94 or higher for all three datasets. One key highlight of *linc2function* from the benchmarking results is that AUC values of *linc2function* have not dropped while used for predicting other species, unlike the other two tools. Essentially, we endeavored to provide a solution that successfully predicts the coding ability of a wide range of species including non-model ones and the results reflect our model’s ability to do so.

**Fig. 10.**
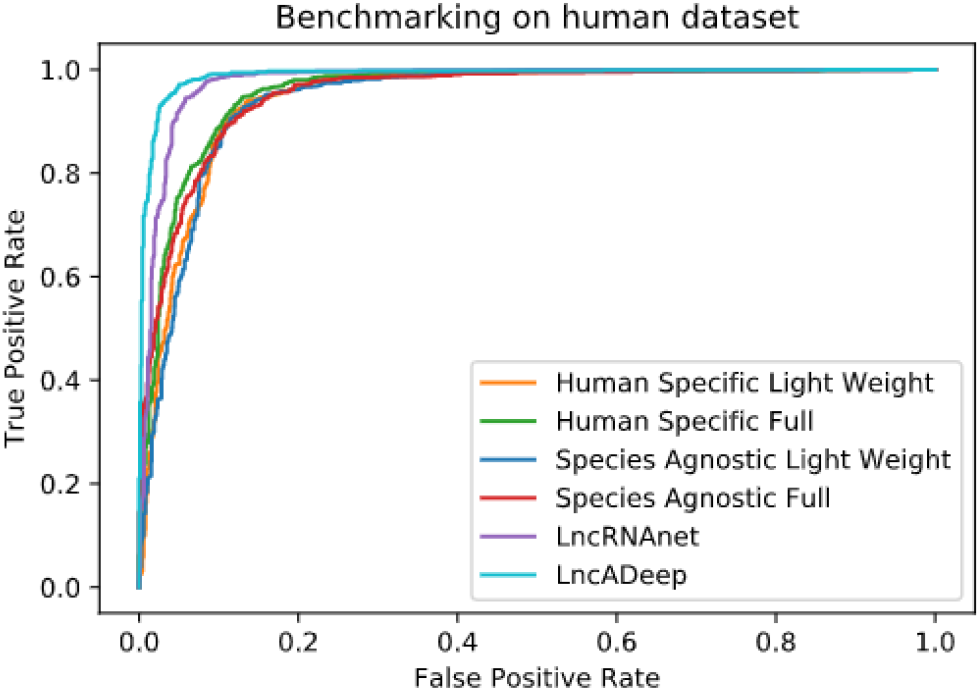
ROC curve for LncADeep, LncRNAnet, linc2function Human Specific, and linc2function Species Agnostic models evaluated on randomly selected Human transcripts from Gencode.

**Fig. 11.**
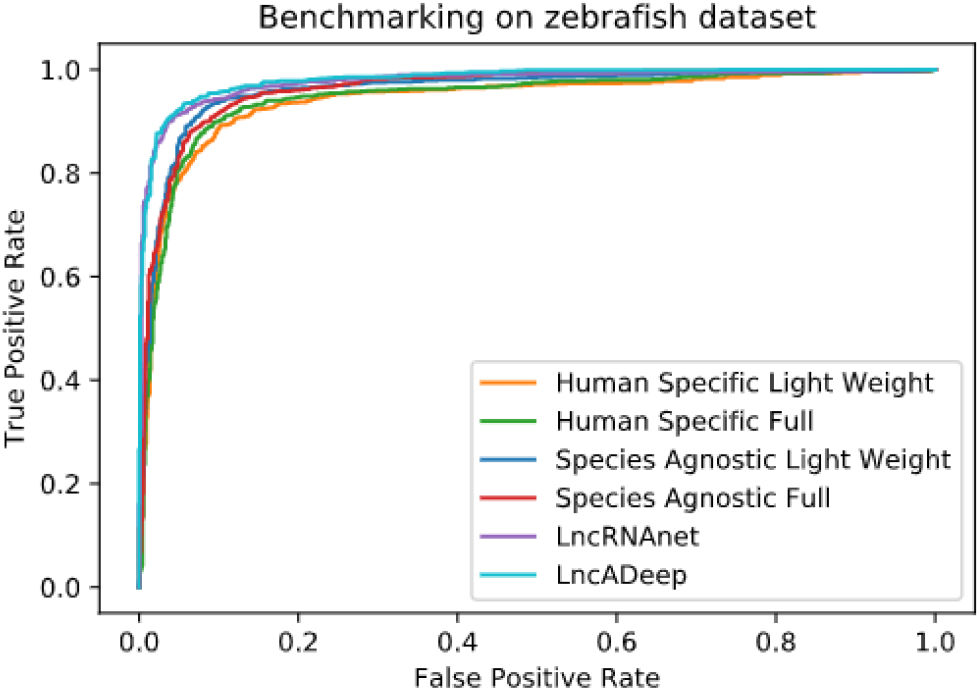
ROC curve for LncADeep, LncRNAnet, linc2function Human Specific, and linc2function Species Agnostic models evaluated on randomly selected Zebrafish transcripts from Ensembl.

**Fig. 12.**
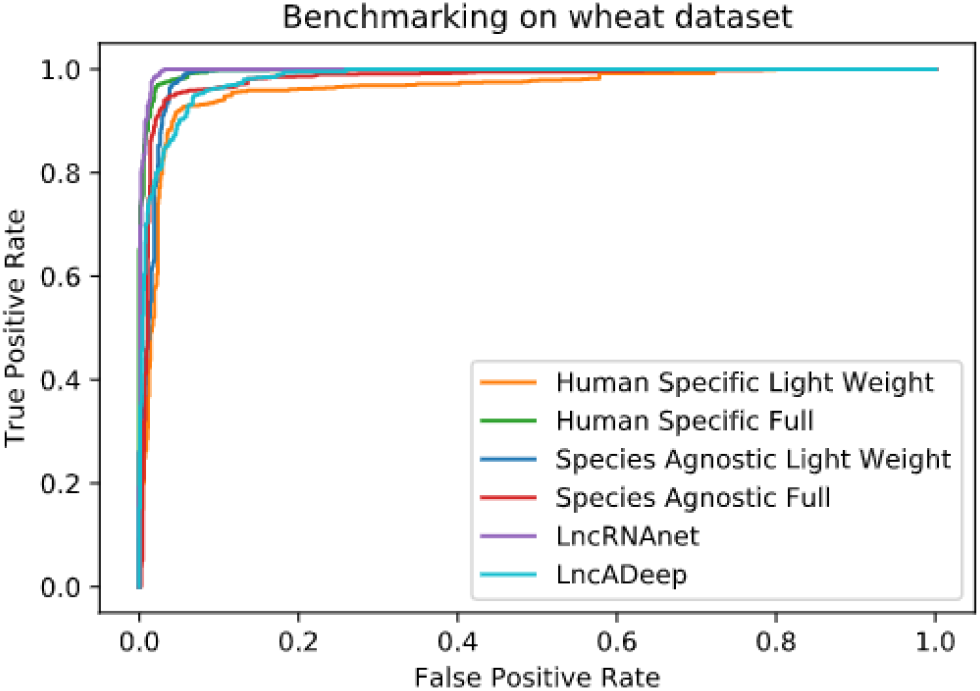
ROC curve for LncADeep, LncRNAnet, linc2function Human Specific and linc2function Species Agnostic models evaluated on randomly selected Wheat transcripts from Ensembl.

**Fig. 13.**
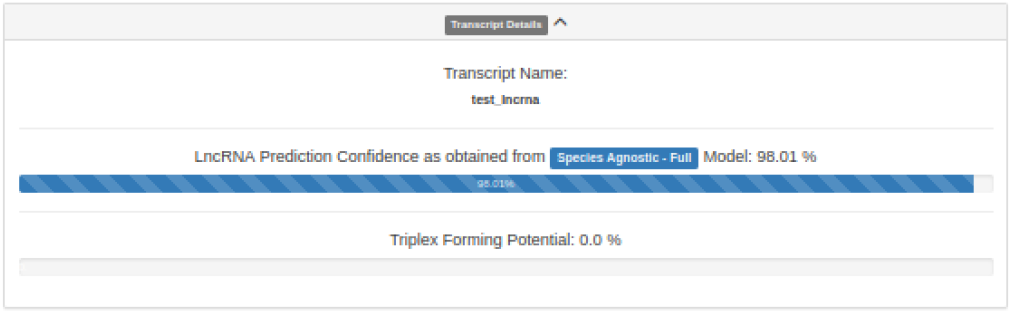
Figure showing transcript details section from linc2function pipeline, displaying its lncRNA forming potential and triplex forming potential (TFP).

**Fig. 14.**
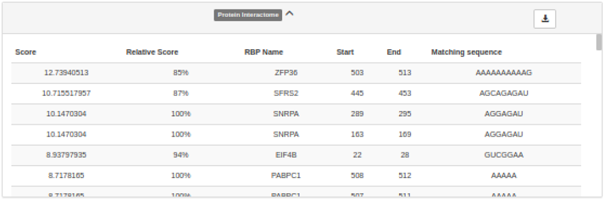
Figure showing protein interactome section from linc2function pipeline, displaying most likely RBP interactions.

**Fig. 15.**
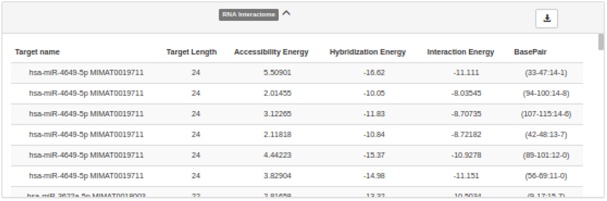
Figure showing RNA interactome section from linc2function pipeline, displaying most likely miRNA interactions.

### 3.2 The *linc2function* Webserver

The webserver can take a FASTA sequence as input and results are displayed as html output along with downloadable text, table and images formats. The first section of the *linc2function* results provides basic details of the predicted transcript such as, the name and length of the input FASTA sequence. Additionally, it contains the lncRNA prediction confidence percentage as predicted from the model selected by the user. Finally, the triplex forming potential of the given transcript is also presented in this section.

The subsequent section contains protein interactome information for the given transcript sequence. It contains interaction score, relative score, RBP name, start position, end position, and matching sequence data of RNA-RBP interactions for the selected list of RNA binding proteins.

The RNA-RBP interactome is followed by the RNA-RNA interactome section which lists the potential miRNA interactions with the lncRNA provided. Details provided in this section include target miRNA name, length of the miRNA target, accessibility/hybridization/interaction energies of the interaction, and bases involved in interaction from the lncRNA and miRNA sequences.

Next two sections shows the predicted secondary structure of the given sequence. Fig 16 shows an example of arc diagram and Fig 17 shows an example of 2D diagram for a representative sequence.

**Fig. 16.**
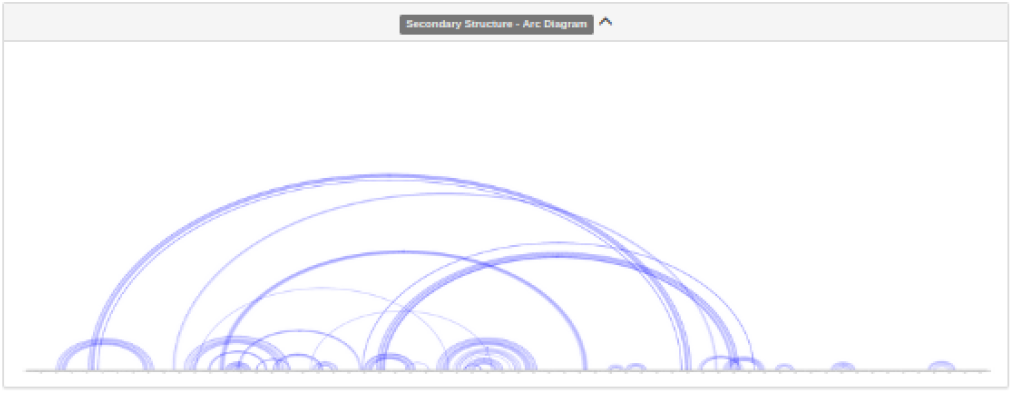
Figure showing secondary structure - arc diagram section from linc2function pipeline, displaying secondary structure in the form of an arc-diagram.

**Fig. 17.**
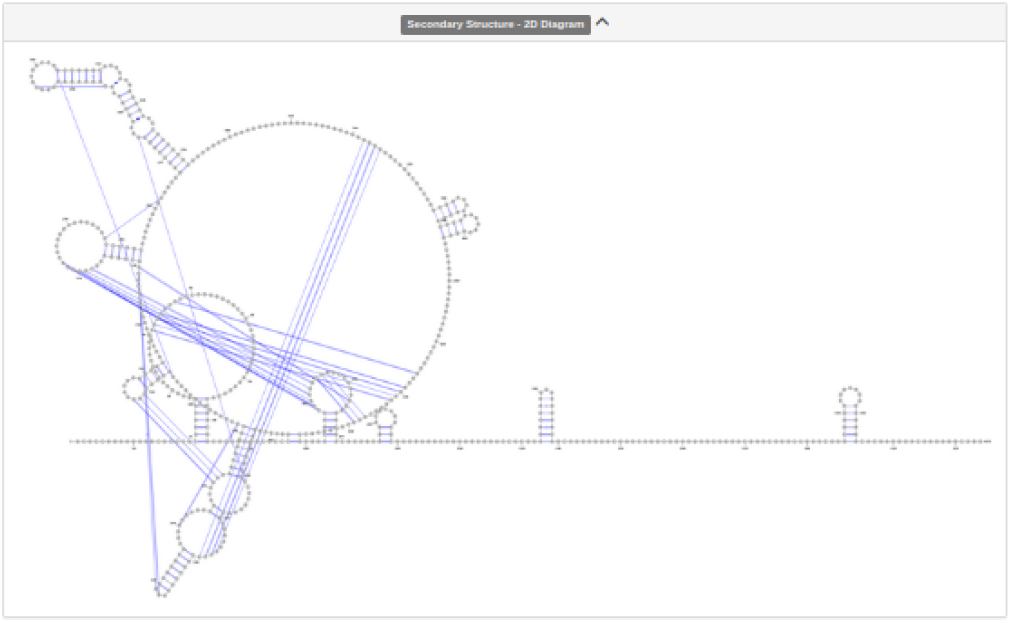
Figure showing secondary structure - 2D diagram section from linc2function pipeline, displaying secondary structure in the form of a 2D diagram.

## 4 Conclusion

Here, we present a comprehensive pipeline called *Linc2function* for identifying and annotating long noncoding RNAs. The pipeline can be run both in a species-specific or species-agnostic manner. Further, by selecting light weight models, the pipeline can also be run in fast mode when screening a large amount of nucleotide data. This flexibility allows user to run our pipeline both on a basic compute infrastructure and high performance computing cluster. The pipeline can be accessed through a webpage and as a standalone installation covering a range of users from bioinformaticians to biologists. The cross-species generalization ability is unique to our approach, which makes it applicable to species that do not have a well annotated reference genome. The pipeline include an ANN model to identify lncRNA transcripts that achieved consistent high AUC values of over 0.9 while testing for 8 disparate species. Our model is flexible and scalable to test newer features or training on new species and open-access code makes it reusable by others in the community. Thus, we present a first end-to-end solution to identify and annotate new lncRNA transcripts involved in disease and development processes.

The source code is made available under MIT license at our GitLab page https://gitlab.com/tyagilab/linc2functionpipeline.

## Author’s contributions

Conceptualisation, S. T Formal analysis, S. T, Y. R, L. K; Funding Acquisition, S. T; Investigation, S. T, Y. R, L. K; Resources, S. T; Supervision, S. T, L. K Visualisation Y. R; Writing – review and editing, S. T, L. K, Y. R.

## Acknowledgements

We thank Tyrone Chen for proofreading the manuscript and providing feedback.

Thanks also to the HPC and MASSIVE (https://www.massive.org.au/) team at the Monash eResearch Platform for providing and supporting this powerful computing cluster.

We also would like to acknowledge and immensely thankful to the open research community behind TriplexFPP [38], RBPDB [39], RIblast [40], and SPOT-RNA [43] for providing the open source tools.

