## Appendix for "*linc2function*: A deep learning model to identify and assign function to long noncoding RNA (lncRNA)"

picture(0,0)(-35,0)(1,0)30 (0,35)(0,-1)30 picture

picture(0,0)(35,0)(-1,0)30 (0,35)(0,-1)30 picture



### A Appendix

Table A1. Classification Report for Human Specific Light Weight Model)

| - | Precision | Recall | F1-Score |
|---|---|---|---|
| Coding | 0.92 | 0.88 | 0.90 |
| Noncoding | 0.88 | 0.91 | 0.89 |
| accuracy | | | 0.90 |
| macro avg | 0.90 | 0.90 | 0.90 |
| weighted avg | 0.90 | 0.90 | 0.90 |

Table A2. Confusion Matrix for Human Specific Light Weight Model)

| - | Positive | Negative |
|---|---|---|
| Positive | 885 | 117 |
| Negative | 81 | 819 |

Positive class: lncRNA, Negative class: other RNA

Table A3. Classification Report for Human Specific Full Model)

| - | Precision | Recall | F1-Score |
|---|---|---|---|
| Coding | 0.92 | 0.89 | 0.90 |
| Noncoding | 0.88 | 0.91 | 0.90 |
| accuracy | | | 0.90 |
| macro avg | 0.90 | 0.90 | 0.90 |
| weighted avg | 0.90 | 0.90 | 0.90 |

Table A4. Confusion Matrix for Human Specific Full Model)

| - | Positive | Negative |
|---|---|---|
| Positive | 890 | 112 |
| Negative | 77 | 823 |

Positive class: lncRNA, Negative class: other RNA

Table A5. Classification Report for Species Agnostic Light Weight Model)

| - | Precision | Recall | F1-Score |
|---|---|---|---|
| Coding | 0.91 | 0.89 | 0.90 |
| Noncoding | 0.88 | 0.91 | 0.89 |
| accuracy | | | 0.89 |
| macro avg | 0.89 | 0.90 | 0.89 |
| weighted avg | 0.90 | 0.89 | 0.89 |

Table A6. Confusion Matrix for Species Agnostic Light Weight Model)

| - | Positive | Negative |
|---|---|---|
| Positive | 887 | 115 |
| Negative | 85 | 815 |

Positive class: lncRNA, Negative class: other RNA

Table A7. Classification Report for Species Agnostic Full Model)

| - | Precision | Recall | F1-Score |
|---|---|---|---|
| Coding | 0.91 | 0.88 | 0.89 |
| Noncoding | 0.87 | 0.90 | 0.89 |
| accuracy | | | 0.89 |
| macro avg | 0.89 | 0.89 | 0.89 |
| weighted avg | 0.90 | 0.89 | 0.89 |

Table A8. Confusion Matrix for Species Agnostic Full Model)

| - | Positive | Negative |
|---|---|---|
| Positive | 887 | 115 |
| Negative | 85 | 815 |

Positive class: lncRNA, Negative class: other RNA

picture(0,0)(-35,0)(1,0)30 (0,-35)(0,1)30 picture

picture(0,0)(35,0)(-1,0)30 (0,-35)(0,1)30 picture

"output" — 2021/1/28 — page 10 — #10

picture(0,0)(-35,0)(1,0)30 (0,35)(0,-1)30 picture picture(0,0)(35,0)(-1,0)30 (0,35)(0,-1)30 picture



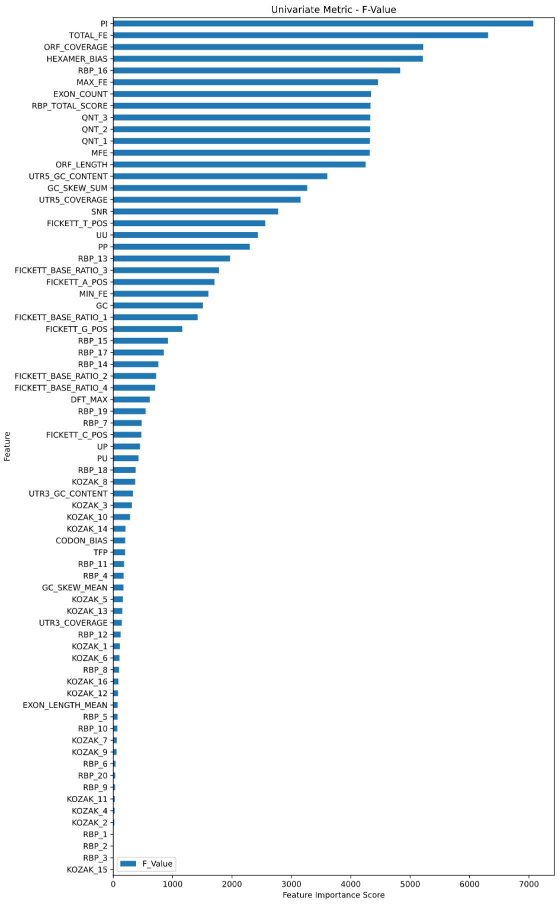


**Fig. A1.** Feature importance measures using F-Value.

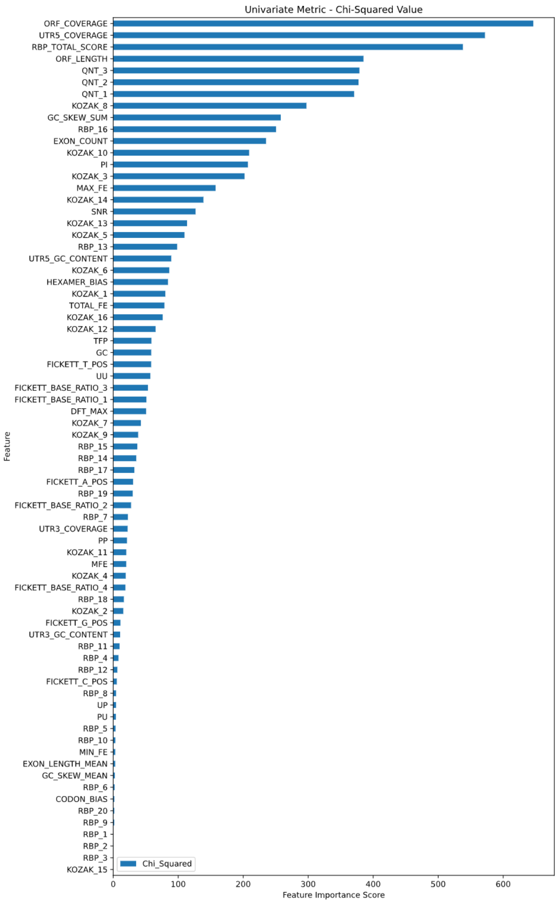


**Fig. A2.** Feature importance measures using Chi-Squared.

picture(0,0)(-35,0)(1,0)30 (0,-35)(0,1)30 picture picture(0,0)(35,0)(-1,0)30 (0,-35)(0,1)30 picture

picture(0,0)(-35,0)(1,0)30 (0,35)(0,-1)30 picture

picture(0,0)(35,0)(-1,0)30 (0,35)(0,-1)30 picture



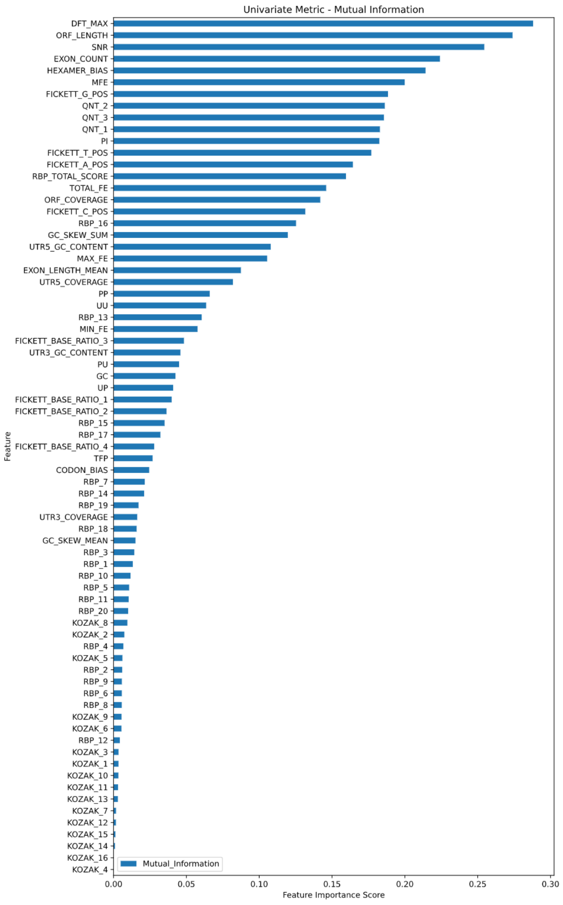


**Fig. A3.** Feature importance measures using Mutual Information.

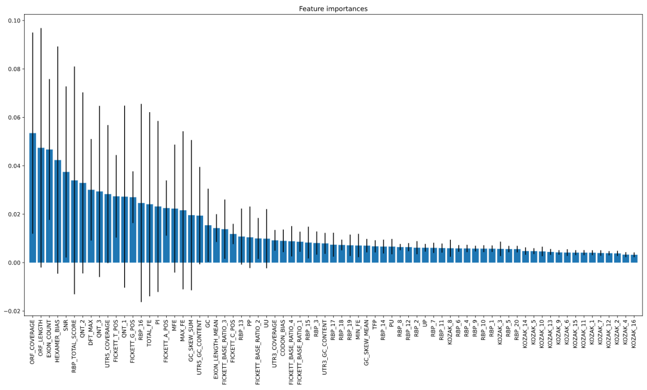


**Fig. A4.** Feature importance measures using tree based models.

picture(0,0)(-35,0)(1,0)30 (0,-35)(0,1)30 picture

picture(0,0)(35,0)(-1,0)30 (0,-35)(0,1)30 picture

“output” — 2021/1/28 — page 12 — #12
